# Complementary role of mathematical modeling in preclinical glioblastoma: differentiating poor drug delivery from drug insensitivity

**DOI:** 10.1101/2021.12.07.471540

**Authors:** Javier C. Urcuyo, Susan Christine Massey, Andrea Hawkins-Daarud, Bianca-Maria Marin, Danielle M. Burgenske, Jann N. Sarkaria, Kristin R. Swanson

## Abstract

Glioblastoma is the most malignant primary brain tumor with significant heterogeneity and a limited number of effective therapeutic options. Many investigational targeted therapies have failed in clinical trials, but it remains unclear if this results from insensitivity to therapy or poor drug delivery across the blood-brain barrier. Using well-established EGFR-amplified patient-derived xenograft (PDX) cell lines, we investigated this question using an EGFR-directed therapy. With only bioluminescence imaging, we used a mathematical model to quantify the heterogeneous treatment response across the three PDX lines (GBM6, GBM12, GBM39). Our model estimated the primary cause of intracranial treatment response for each of the lines, and these findings were validated with parallel experimental efforts. This mathematical modeling approach can be used as a useful complementary tool that can be widely applied to many more PDX lines. This has the potential to further inform experimental efforts and reduce the cost and time necessary to make experimental conclusions.

**Author summary:** Glioblastoma is a deadly brain cancer that is difficult to treat. New therapies often fail to surpass the current standard of care during clinical trials. This can be attributed to both the vast heterogeneity of the disease and the blood-brain barrier, which may or may not be disrupted in various regions of tumors. Thus, while some cancer cells may develop insensitivity in the presence of a drug due to heterogeneity, other tumor areas are simply not exposed to the drug. Being able to understand to what extent each of these is driving clinical trial results in individuals may be key to advancing novel therapies. To address this challenge, we used mathematical modeling to study the differences between three patient-derived tumors in mice. With our unique approach, we identified the reason for treatment failure in each patient tumor. These results were validated through rigorous and time-consuming experiments, but our mathematical modeling approach allows for a cheaper, quicker, and widely applicable way to come to similar conclusions.

## Introduction

Glioblastoma (GBM) is the most common primary brain malignancy and is aggressive, heterogeneous, and diffusely-invasive. Despite maximal surgical resection, radiation therapy, and chemotherapy, patients with GBM have a two-year survival of approximately 25%, and five-year survival under 10% [1–3]. In efforts to further improve overall survival of patients with GBM, many targeted therapies and other chemotherapeutic drugs have been developed and appear promising in preclinical trials [4]. Yet, many of these drugs fail at the clinical level. To this day, none have surpassed the standard of care established over a decade ago, leaving the median overall survival at a dismal 14.6 months [2].

Drug delivery to the brain is an inherent challenge posed by the blood-brain barrier (BBB), which largely restricts hydrophilic molecules and large macromolecules in the bloodstream from crossing into the brain [5]. While this is a critical aspect of protecting this vital organ against various infections, it also prevents a vast majority of oral or intravenous drugs from being delivered to brain tumors. This inherent limitation in drug delivery is always a consideration when investigating the cause of targeted therapies that failed to show a clinical benefit. Exacerbating this challenge, GBM is notorious for its inter- and intra-tumoral heterogeneity. Beyond questions of drug delivery, researchers must also consider the presence of drug-resistant tumor cells, which varies on a per patient basis. This further limits the interpretation of clinical trials, which traditionally focus on the average outcomes of populations. If researchers could effectively differentiate the cause of targeted drug therapy failure in clinical trials on a per patient basis, it would undoubtedly result in more approved targeted drug options available for patients.

However, as it is not easy to determine where drug was actually delivered in a human brain, the gold standard for investigating drug failure is via *in vivo* experimentation with cell lines and animal models. To form definitive conclusions, numerous repeats are required, with tumors grown outside of and within the brain, and the resulting tissue undergoing DNA, RNA, and protein sequence analyses. Recently, Marin *et al*. did just this for depatuxizumab mafodotin (Depatux-M, ABT-414), an EGFR-directed antibody drug conjugate (ADC) [6]. In brief, they performed intracranial experiments for seven different cell lines, and through numerous experiments with each line, they determined that only two cell lines were responsive to the therapy. Of the remaining five cell lines, two acquired resistance to the drug and three were not responsive to the drug due to a relatively intact BBB. This work was the first of its kind to go so deep and to fully highlight the challenges facing the development of targeted drugs.

Recently, in Massey *et al*., we detailed our approach of how a mathematical model could utilize only a subset of the data collected in Marin *et al*. to identify the driving factor of drug failure: limited drug delivery or the presence of a population insensitive to treatment [6, 7]. This is significant because it enables researchers to quickly assess cell lines and determine which deserve deeper experimentation. By reducing time and costs, we hope that this could focus drug development research and increase the speed of acquiring clinically meaningful answers about drug effectiveness.

In this paper, we demonstrate that our model, using relatively minimal experimental data, arrives at similar conclusions to the highly sophisticated experiments conducted in Marin *et al*. for a subset of the cell lines [6]. First, we describe the mathematical model as presented in Massey *et al*. [7]. Then, we provide a brief description of the cell lines and experiments that were used. We briefly discuss the methodology for parameterization and present our findings for each of the cell lines. We then conclude with a few thoughts regarding the implications of our model and its hopeful future use.

## Methods

### Preclinical Experiment: Data Collection

Three cell lines were selected from The Mayo Clinic Brain Tumor Patient-Derived Xenograft (PDX) National Resource [8] for their known differential response to Depatux-M (see S1 Table for further details). These cell lines were transduced with firefly luciferase (F-luc) to allow for bioluminesence imaging (BLI). As the resulting BLI flux is linearly proportional to the total number of luminescing cells, we can non-invasively monitor tumor progression *in vivo* over time [9]. In brief, patient-derived GBM cells were implanted heterotopically into ten athymic nude mice per cell line.

After the tumors were established, PDXs were assigned to therapy groups, including a sham control and Depatux-M. Therapy was initiated a set number of days post injection (7d, 14d, or 21d) and subsequently administered on 7 day intervals. To explore response under the effect of the BBB, a similar experiment followed PDXs with tumors implanted intracranially. BLI was acquired 1-3 times weekly, with less frequent imaging after 4 weeks post-initiation. All animal studies were approved by the Mayo Institutional Animal Care and Use Committee. Per IACUC guidelines, upon reaching a moribund state, PDX mice were euthanized and data collection ceased. The collected bioluminescence data, as collected in Marin *et al*., is shown in Fig 1 [6].

**Fig 1.**
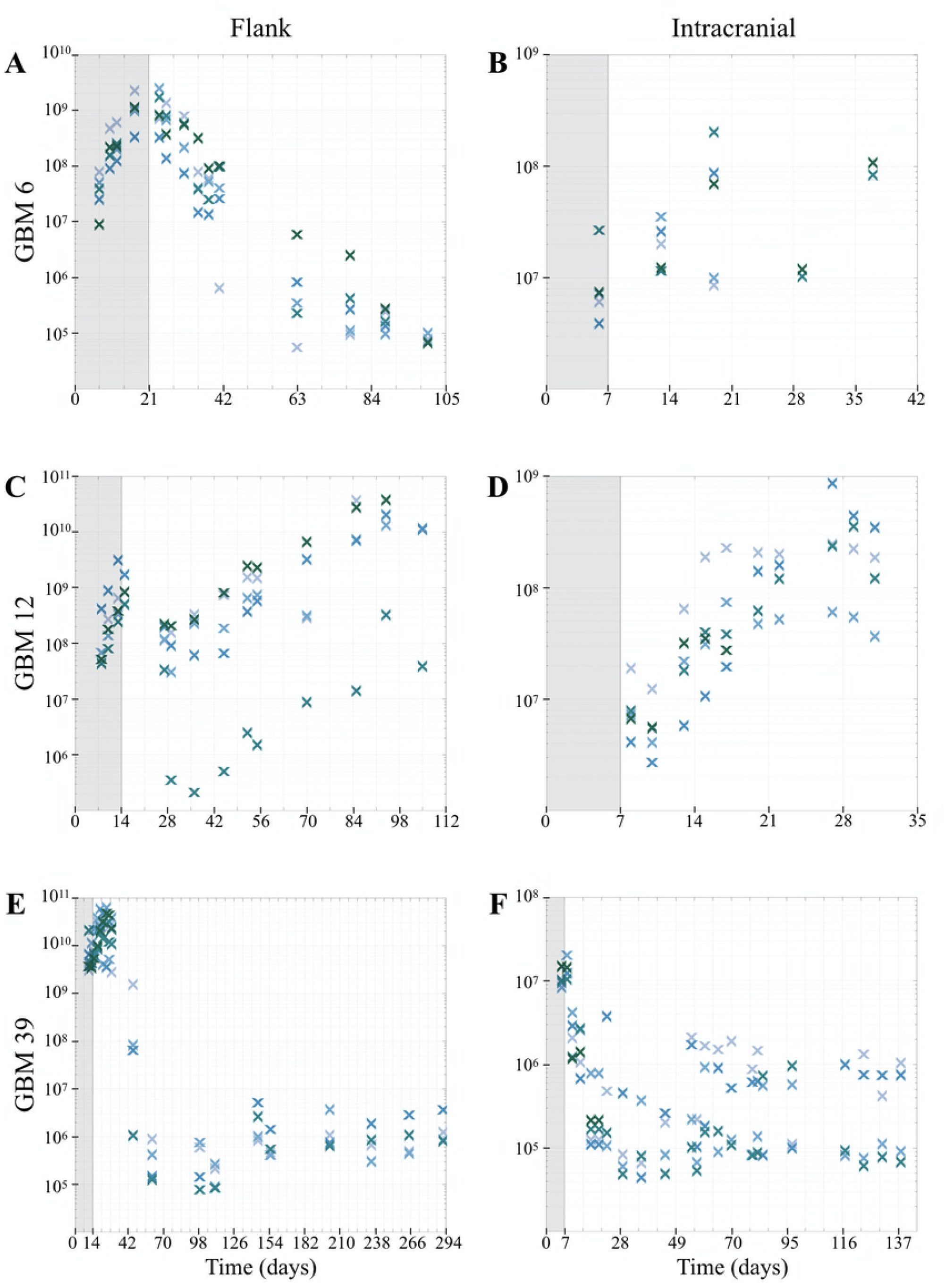
Experimental data for PDXs treated with the ADC. BLI flux data was recorded in photons/sec. Shaded area indicates time prior to treatment initiation. Five subjects were utilized in each experiment, as denoted a-f: (**a**) GBM6, flank; (**b**) GBM6, intracranial; (**c**) GBM12, flank; (**d**) GBM12, intracranial; (**e**) GBM39, flank; (**f**) GBM39, intracranial.

As expected, tumors treated with a sham control grew exponentially. Heterogeneous responses to treatment were noted across the selected cell lines, paired with additional variability amongst mice of the same cell line. In GBM6, tumor size decreased in the flank environment, but appeared to maintain size intracranially, despite treatment. In GBM12, flank tumors had a modest response to the therapy, shortly followed by regrowth. Nevertheless, intracranial GBM12 tumors did not respond to therapy. On the other hand, GBM39 tumors responded very well to treatment in both the flank and intracranially.

This varying success of Depatux-M *in vivo* can likely be attributed to two major factors, both of which were captured with the preclinical experimental design. The flank data captured tumor sensitivity to drug, and the intracranial data incorporated the additional aspect of the BBB. Particularly, we take a two-pronged approach that uses this experimental data to investigate the relative contributions that ADC insensitivity and BBB permeability have on drug failure, leading to persistent tumor growth.

### Mathematical Model: Treatment Exposure and Sensitivity

In this paper, we utilize the Treatment Exposure and Sensitivity model, first presented in Massey *et al*. [7]. It was designed to differentiate the relative contributions of cell sensitivity and drug delivery to the overall tumor response to therapy. We used the previously described preclinical experimental data to parameterize this model.

### Equations and Parameters

The model is a system of three ordinary differential equations (1) capturing the dynamics of the drug (*A*) and two underlying tumor populations: one subpopulation that is highly sensitive to the treatment (*H*) and another that is less sensitive (*L*). The model, as listed below,

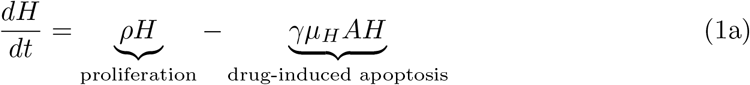

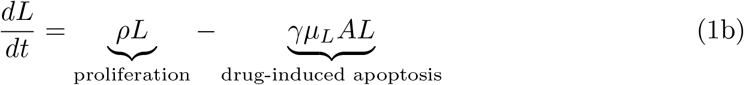

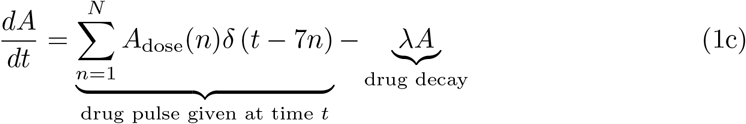

has the analytical solution

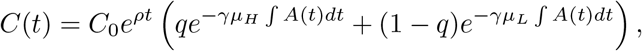

where *C*(*t*) = *H*(*t*) + *L*(*t*), and

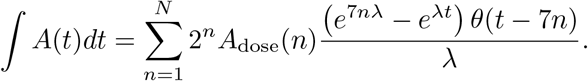

Parameters for the model are summarized in Table 1.

**Table 1.**
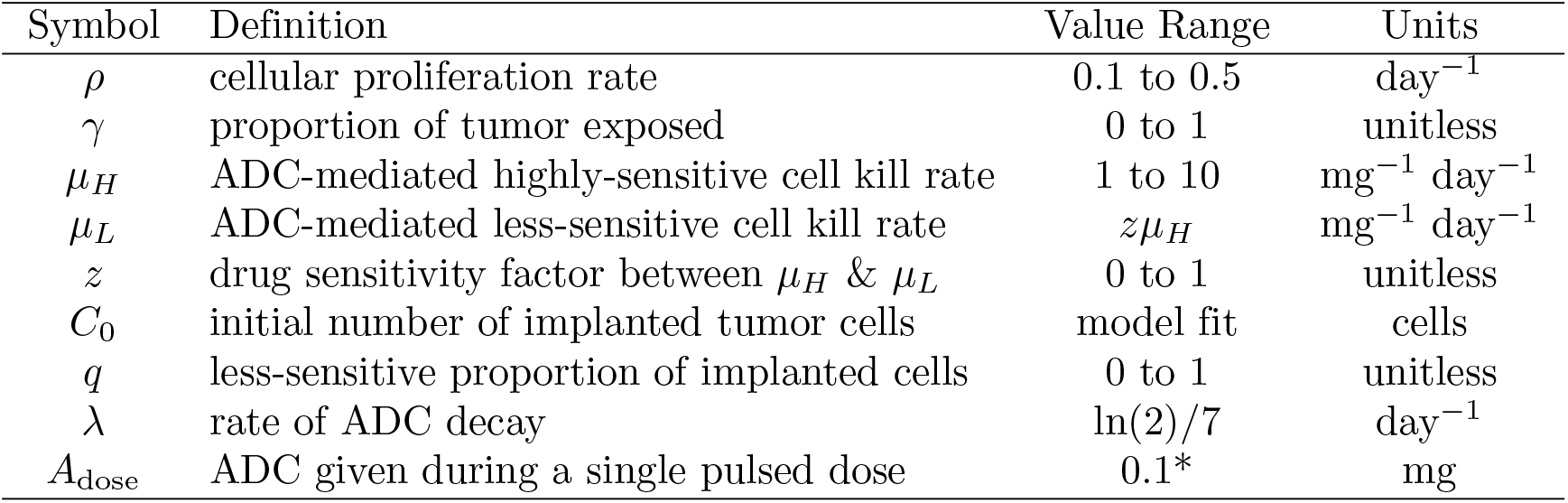
Model Parameter Definitions and Values. Adapted from Massey *et al*. [7]. *Note: ADC dose was given according to animal weights, but variation was unremarkable so we modeled the dose as a constant.

### Parameter Estimation

Distinct from Massey *et al*., a Bayesian approach was employed for fitting parameters in an attempt to capture the generic behavior of the cell line while acknowledging the varying behavior in individual mice [7]. Similar to Massey *et al*. though, parameters were calibrated in a sequential process enabled by the experimental design [7].

### Estimating proliferation rate (*ρ*)

The calibration of the proliferation rate *ρ* was done in two parts for both the flank and intracranial settings. First, as the sham control experiments have no therapeutic effect, the model can be evaluated without treatment, thereby simplifying to an exponential growth model. Using the Matlab **®** (MATLAB Release 2018b, The MathWorks, Inc., Natick, Massachusetts, United States) `lsqcurvefit` function, the logged-data can be used to fit the initial number of viable tumor cells and the proliferation rate *ρ*. Since proliferation rate is both intrinsic to the PDX line and specific to the microenvironment, the subject-specific fits obtained in fitting the sham control data was used to develop an informative Gaussian prior. This was then used to estimate the proliferation rate for treatment in the same microenvironment.

### Estimating sensitivity rates (*µ*_*H*_, *µ*_*L*_), the insensitive tumor proportion (*q*), and proportion of drug delivered (*γ*)

The calibration of the sensitive and insensitive proportion *q* and respective rates *µ*_*H*_, *µ*_*L*_ was specific to the flank or intracranial settings. Since the flank does not have the limitation of the BBB, we can assume that the entirety of the tumor is exposed to the therapy and that *γ* = 1. The remaining parameters *q, µ*_*H*_, and *µ*_*L*_ were sampled from uninformative priors spanning the ranges in Table 1. The selected parameter set was subject-specific and minimized the sum-of-squared-error for that subject in the flank. With five subjects in the flank, we formulated a Gaussian prior for the *µ*_*H*_ sensitivity using the selected parameter sets. Using this prior, we proceed with a similar approach to select remaining parameter sets in the intracranial setting.

## Results

Applying this fitting algorithm to each of the three PDX lines, variations in the model parameters captured the wide range of treatment response dynamics in the data, both across and within experiments. The resulting fits are displayed in Fig 2, where each color represents a different experimental subject. Values for fitted model parameters are tabulated in S2 Table - S7 Table. Of particular interest, parameter values for drug sensitivity and BBB penetrance are plotted in Fig 3, where GBM6 is represented by red squares, GBM12 is represented by green triangles, and GBM39 is represented by the blue circles.

**Fig 2.**
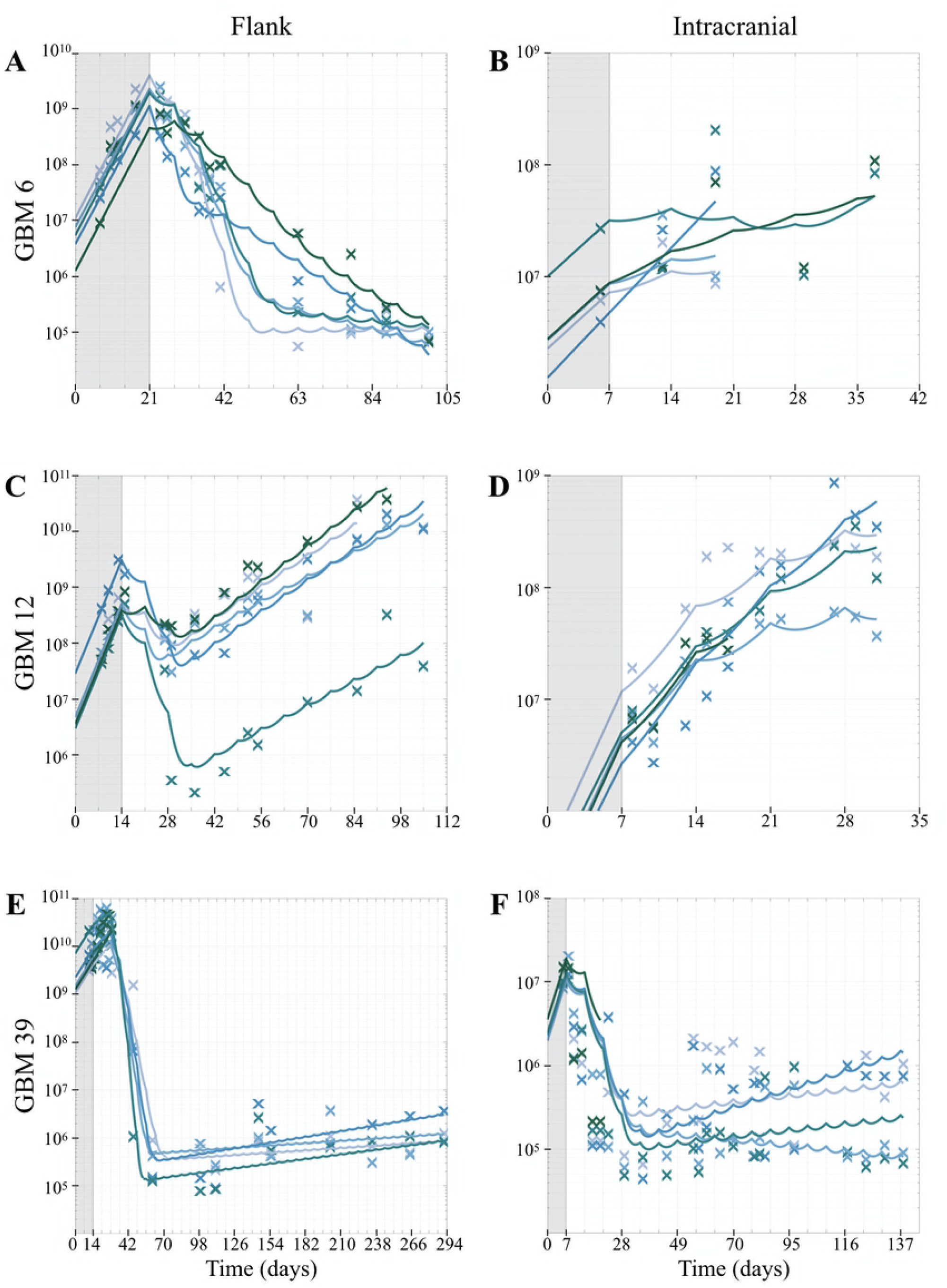
Treatment Exposure and Sensitivity model fits to experimental data. Recall, BLI flux data was recorded in photons/sec and treatment was initiated after the shaded area. Five subjects were randomized to each condition, shown here with their respective fits. The fitting process was repeated for sets of data, denoted A-F: (**a**) GBM6, flank; (**b**) GBM6, intracranial; (**c**) GBM12, flank; (**d**) GBM12, intracranial; (**e**) GBM39, flank; (**f**) GBM39, intracranial.

**Fig 3.**
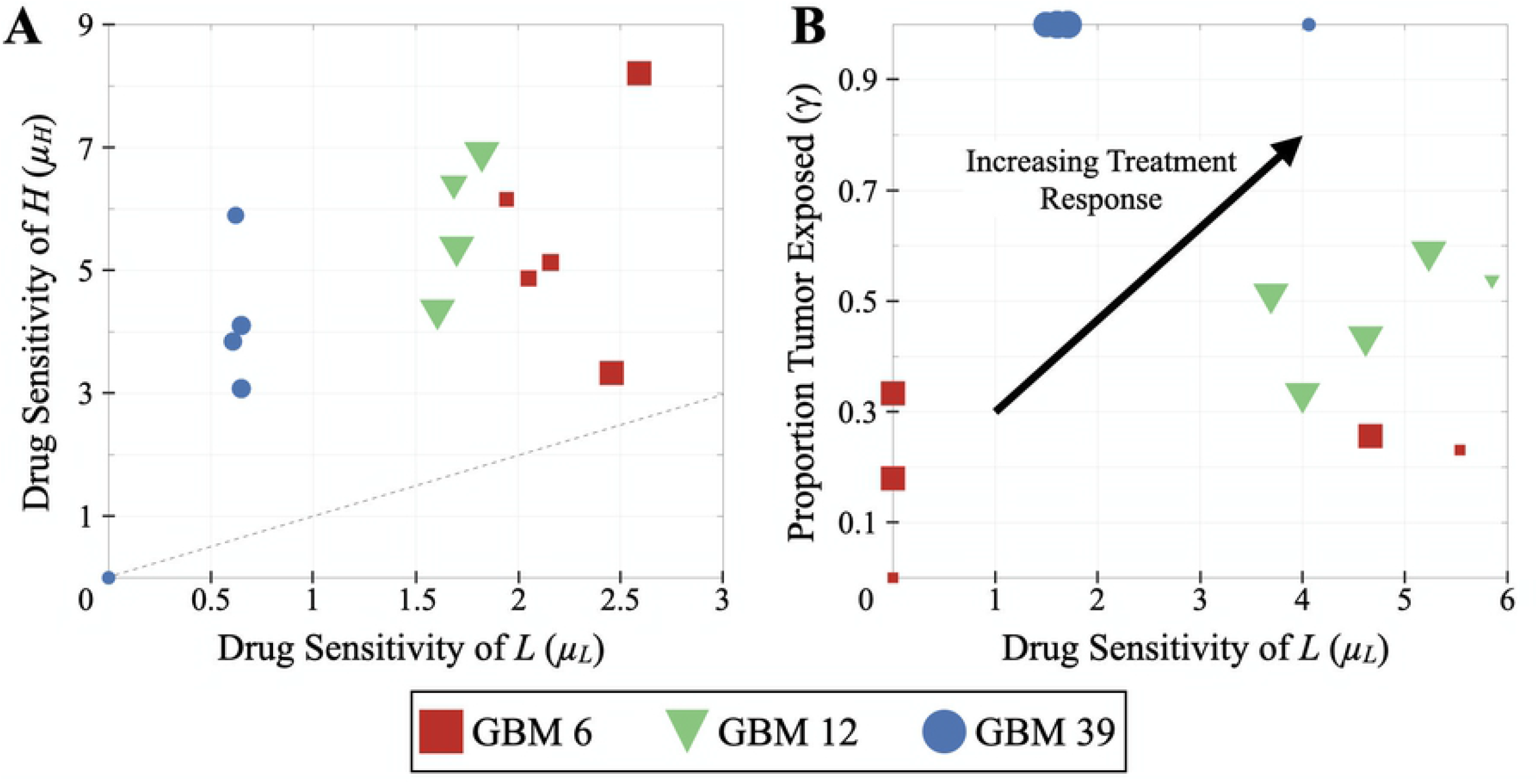
Parameter values resulting from the fitted experimental data. Marker size corresponds to the magnitude of *q*, the estimated proportion of less-sensitive cells initially implanted. (**a**) Parameter estimates for the drug sensitivity of L and H (*µ*_*L*_ and *µ*_*H*_, respectively) in the flank setting. The dotted line represents the line of identity, where *µ*_*L*_ = *µ*_*H*_; (**b**) Parameter estimates for the drug sensitivity of L (*µ*_*L*_) is plotted against the proportion of tumor exposed (*γ*) in the intracranial setting.

### Highly-sensitive rate (*µ*_*H*_) consistent across all PDX lines

The highly-sensitive cell kill rate (*µ*_*H*_) was consistent (3.1–8.2 mg^*−*1^day^*−*1^, excluding one GBM39 subject that failed to respond) for all flank estimates in PDX lines, as shown in Fig 3A. This suggests that the highly-sensitive population *H* within each PDX line are fairly similar. When applying the Gaussian distribution for the intracranial experiments, the parameter estimations remained in this range. Both GBM6 and GBM12 were estimated to have similar *µ*_*H*_ values (5.5 and 5.8 mg^*−*1^day^*−*1^, respectively), and the highly-sensitive population *H* of GBM39 was estimated to be slightly less sensitive than the other PDX lines (average of 4.2 mg^*−*1^day^*−*1^).

### Less-sensitive kill rate (*µ*_*L*_) varies by PDX line

In contrast, estimates of the less-sensitive cell kill rate (*µ*_*L*_) captured the differences between PDX lines and was consistent across subjects. In the flank experiments, the average rates for GBM6, GBM12, and GBM39 were 2.2, 1.7, and 0.6 mg^*−*1^day^*−*1^, respectively (Fig 3A). This suggests that, of the PDX lines investigated, the less-sensitive population of GBM6 was the most sensitive and GBM 39 the least sensitive in the flank. Intracranially, this trend was much more obscure. On average, the estimated intracranial *µ*_*L*_ values for GBM6, GBM12, and GBM39 were 5.1 (excluding three GBM6 subjects that failed to respond), 4.7, and 2.1 mg^*−*1^day^*−*1^ (Fig 3B). This suggests that the less-sensitive population *L* of GBM39 appears to be less sensitive than the GBM6 and GBM12 intracranially.

### Proportion of tumor exposed to drug (*γ*) varies by PDX line

Estimates of the proportion of tumor exposed to the drug (*γ*) also captured the variability between PDX lines. Recall, this parameter was set to 100% for the flank experiments. Intracranially, the model estimated that, on average, 20% of drug was successfully delivered for GBM6, 48% for GBM12, and 100% for GBM39 (Fig 3B). While GBM6 appeared to be the most sensitive of the PDX lines, the poor drug delivery limited the response intracranially. Similarly, although GBM39 was estimated to be the least sensitive of the PDX lines, the estimated high drug delivery resulted in a favorable response to intracranial therapy. This suggests that drug delivery plays a major role in treatment response.

## Discussion

Despite numerous clinical trials investigating the use of novel and repurposed therapies for GBM, the standard of care has not changed since 2005. The influence of the BBB is a key area of study, as it poses a significant confounding factor by limiting drug delivery. As a result, it remains unclear if cells are inherently resistant to these therapies or if it is simply not reaching its target. By identifying the root cause of these failed clinical trials, we can better inform researchers and clinicians in evaluating the benefits of particular drugs.

This work builds upon our effort in Massey *et al*., where the model was first presented, by demonstrating the utility of the model through applying the parameter estimation to serial BLI data from three PDX lines [7]. Broadly, we found the model was able to fit data from multiple replicates of multiple cell lines. Through the key parameter fittings of *µ*_*L*_ and *γ*, the model provides insights into the different underlying mechanisms driving the tumor growth behavior after therapy in the intracranial setting. For GBM6, the model attributed the intracranial treatment failure to poor drug delivery, presumably due to an intact BBB. While GBM12 was also fairly sensitive to the drug, the model identified poor delivery as the major limitation, again presumably as a result of the BBB. Unlike the other PDX lines, GBM39 responded well to treatment. The model primarily attributed this to high drug delivery, suggestive of a permeable BBB.

By using only serial BLI data, the model was able to make similar inferences about the three PDX cell lines as the more rigorous results found in Marin *et al*. [6]. For example, the model identified drug delivery as a key contributor to the outcomes of GBM6 and GBM39. In fixed brain sections, elevated levels of fibrinogen are indicative of BBB disruption and while GBM6 demonstrated no detectable fibrinogen accumulation, GBM39 tumors had significant fibrinogen buildup near the region of the tumor. Further, GBM6 tumors demonstrated minimal accumulation of drug while GBM39 tumors exhibited drug accumulation near the region of the tumor [6].

The agreement, though not perfect, between these two approaches is very encouraging. The BLI experiments used in this paper were conducted in parallel to the experiments for Marin *et al*. [6]. While it was decided to publish Marin *et al*. first, the results presented here were identified first [6]. We believe this demonstrates the promising complementary utility our model brings to investigating drug failures. BLI experiments are relatively quick and cheap compared to the many rigorous experiments performed in Marin *et al*. [6]. Our model can be easily applied to additional PDX lines, and, in future investigations, could be used to identify which lines warrant more detailed experiments. For example, it may identify which lines are anticipated to have the most similar or different underlying response mechanisms. We believe this coupled, complementary modeling and experimental approach could lead to a more efficient pipeline for understanding drug responses.

## Conclusion

This work has demonstrated how the use of a mathematical model with relatively minimal experimental data can be used to predict the primary factor that contributes to intracranial drug failure. While it should not be expected that this model will replace more complicated experimental methodologies for deep biological understanding of drug failure, as done in Marin *et al*., we believe the model can serve in a complementary fashion, streamlining initial cell line selection for more rigorous experiments [6]. In this way, the model could reduce overall experimental costs and hopefully also decrease time to meaningful answers for patients.

## Supporting information

**S1 Table.** PDX cell lines. Additional details about the primary tumors that the 3 PDXs were derived from.

| PDX Line | Sex | EGFR status | D/rho |
| --- | --- | --- | --- |
| GBM6 | M | Amp, vIII | 4.7 |
| GBM12 | M | Amp, G719A | 1.7 |
| GBM39 | M | Amp, vIII | 0.9 |

**S2 Table.** GBM6 Flank Fit Values. Parameter fits obtained from model fitting. Accuracy was measured by the sum of squared error between the logged data and the logged model simulation (termed ssLe).

| Parameter | Subject 1 | Subject 2 | Subject 3 | Subject 4 | Subject 5 |
| --- | --- | --- | --- | --- | --- |
| $\rho$ | 0.2806 | 0.2715 | 0.2715 | 0.2806 | 0.2806 |
| $\mu_H$ | 5.1538 | 5.1282 | 8.2051 | 4.8718 | 3.3333 |
| $\mu_L$ | 1.6276 | 2.1595 | 2.5912 | 2.0515 | 2.4560 |
| $\gamma$ | 1 | 1 | 1 | 1 | 1 |
| $q$ | -5.3684 | -4.1053 | -2 | -4.5263 | -2 |
| $C_0$ | $1.19 * 10^7$ | $7.59 * 10^6$ | $3.78 * 10^6$ | $5.43 * 10^6$ | $1.26 * 10^6$ |
| ssLe | 10.6708 | 4.5722 | 3.2533 | 7.6001 | 15.8735 |

**S3 Table.** GBM6 Intracranial Fit Values. Parameter fits obtained from model fitting. Accuracy was measured by the sum of squared error between the logged data and the logged model simulation (termed ssLe).

| Parameter | Subject 1 | Subject 2 | Subject 3 | Subject 4 | Subject 5 |
| --- | --- | --- | --- | --- | --- |
| $\rho$ | 0.1652 | 0.1652 | 0.1913 | 0.1652 | 0.1652 |
| $\mu_H$ | 5.5390 | 5.5390 | 5.5390 | 5.5390 | 5.5390 |
| $\mu_L$ | 4.6644 | 5.5390 | 0 | 0 | 0 |
| $\gamma$ | 0.2564 | 0.2308 | 0 | 0.3333 | 0.1795 |
| $q$ | -2 | -10 | -10 | -2 | -2 |
| $C_0$ | $2.26 * 10^6$ | $2.69 * 10^6$ | $1.24 * 10^6$ | $9.97 * 10^6$ | $2.75 * 10^6$ |
| ssLe | 0.5248 | 1.2012 | 0.7040 | 6.0080 | 3.1166 |

**S4 Table.** GBM12 Flank Fit Values. Parameter fits obtained from model fitting. Accuracy was measured by the sum of squared error between the logged data and the logged model simulation (termed ssLe).

| Parameter | Subject 1 | Subject 2 | Subject 3 | Subject 4 | Subject 5 |
| --- | --- | --- | --- | --- | --- |
| $\rho$ | 0.3339 | 0.3339 | 0.3339 | 0.3339 | 0.3339 |
| $\mu_H$ | 5.3846 | 6.9231 | 6.4103 | 6.9231 | 4.3590 |
| $\mu_L$ | 1.7005 | 1.8222 | 1.6872 | 1.8222 | 1.6059 |
| $\gamma$ | 1 | 1 | 1 | 1 | 1 |
| $q$ | -2 | -2 | -3.2632 | -4.1053 | -2 |
| $C_0$ | $4.74 * 10^6$ | $4.76 * 10^6$ | $2.87 * 10^7$ | $3.01 * 10^6$ | $3.57 * 10^6$ |
| ssLe | 10.8783 | 6.2766 | 3.6334 | 14.9513 | 4.0026 |

**S5 Table.**
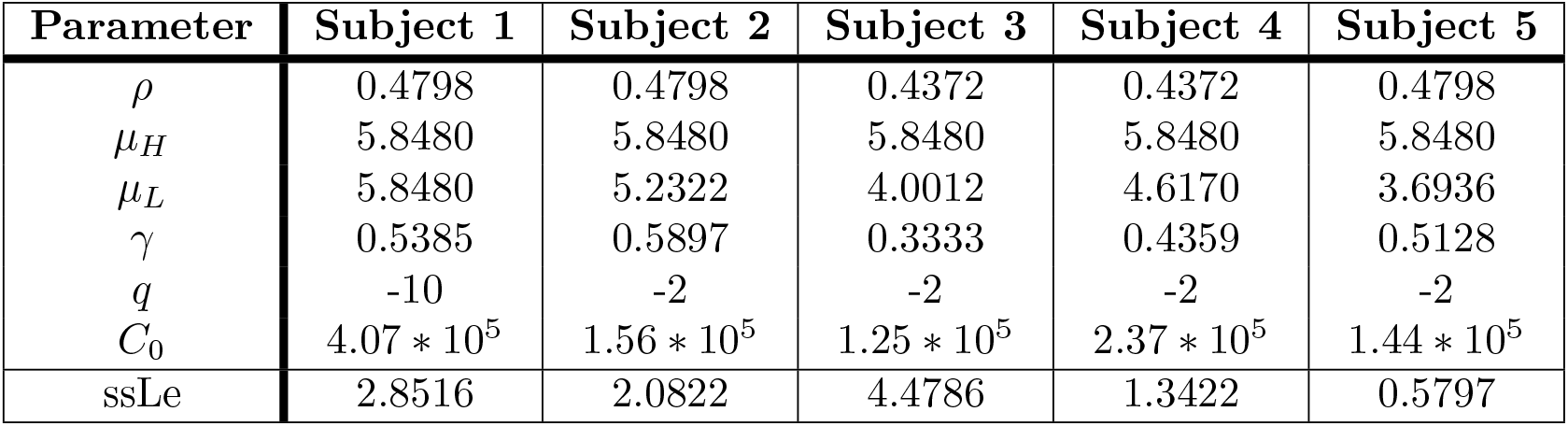
GBM12 Intracranial Fit Values. Parameter fits obtained from model fitting. Accuracy was measured by the sum of squared error between the logged data and the logged model simulation (termed ssLe).

| Parameter | Subject 1 | Subject 2 | Subject 3 | Subject 4 | Subject 5 |
| --- | --- | --- | --- | --- | --- |
| $\rho$ | 0.4798 | 0.4798 | 0.4372 | 0.4372 | 0.4798 |
| $\mu_H$ | 5.8480 | 5.8480 | 5.8480 | 5.8480 | 5.8480 |
| $\mu_L$ | 5.8480 | 5.2322 | 4.0012 | 4.6170 | 3.6936 |
| $\gamma$ | 0.5385 | 0.5897 | 0.3333 | 0.4359 | 0.5128 |
| $q$ | -10 | -2 | -2 | -2 | -2 |
| $C_0$ | $4.07 * 10^5$ | $1.56 * 10^5$ | $1.25 * 10^5$ | $2.37 * 10^5$ | $1.44 * 10^5$ |
| ssLe | 2.8516 | 2.0822 | 4.4786 | 1.3422 | 0.5797 |

**S6 Table.** GBM39 Flank Fit Values. Parameter fits obtained from model fitting. Accuracy was measured by the sum of squared error between the logged data and the logged model simulation (termed ssLe).

| Parameter | Subject 1 | Subject 2 | Subject 3 | Subject 4 | Subject 5 |
| --- | --- | --- | --- | --- | --- |
| $\rho$ | 0.0975 | 0.0975 | 0.0975 | 0.0975 | 0.09075 |
| $\mu_H$ | 3.0769 | 4.1026 | 3.8462 | 5.8974 | 0.0000 |
| $\mu_L$ | 0.6477 | 0.6478 | 0.6073 | 0.6210 | 0.0000 |
| $\gamma$ | 1 | 1 | 1 | 1 | 1 |
| $q$ | -4.9474 | -4.9474 | -5.3684 | -6.2105 | -10 |
| $C_0$ | $1.05 * 10^9$ | $1.48 * 10^9$ | $2.21 * 10^9$ | $7.10 * 10^9$ | $1.25 * 10^9$ |
| ssLe | 6.1545 | 5.8926 | 6.1951 | 5.8559 | 6.9761 |

**S7 Table.** GBM39 Intracranial Fit Values. Parameter fits obtained from model fitting. Accuracy was measured by the sum of squared error between the logged data and the logged model simulation (termed ssLe).

| Parameter | Subject 1 | Subject 2 | Subject 3 | Subject 4 | Subject 5 |
| --- | --- | --- | --- | --- | --- |
| $\rho$ | 0.2394 | 0.2394 | 0.2394 | 0.2394 | 0.2394 |
| $\mu_H$ | 4.3538 | 4.0636 | 4.0636 | 4.3538 | 4.0636 |
| $\mu_L$ | 1.6039 | 1.7112 | 1.4970 | 1.6039 | 4.0636 |
| $\gamma$ | 1 | 1 | 1 | 1 | 1 |
| $q$ | -2.4211 | -2.4211 | -2.8421 | -2.8421 | -10 |
| $C_0$ | $2.34 * 10^6$ | $1.98 * 10^6$ | $2.25 * 10^6$ | $2.40 * 10^6$ | $3.54 * 10^6$ |
| ssLe | 46.0727 | 23.4508 | 45.3879 | 35.2899 | 29.0847 |

## Acknowledgments

The ADC, depatuxizumab mafodotin, was provided by AbbVie.

